# Neural shape mediation analysis

**DOI:** 10.1101/2023.09.21.558808

**Authors:** Jules Brochard, Etienne Combrisson, Benoit Chatard, Jean-Philippe Lachaux, Andrea Brovelli, Jean Daunizeau

**Affiliations:** Inserm U 1127, CNRS UMR 7225, Sorbonne Universités, Institut du Cerveau, ICM, Paris, France; Centre de Recherche en Neurosciences de Lyon (CRNL), Institut National de la Santé et de la Recherche Médicale (INSERM), Lyon, France; Institut de Neurosciences de la Timone, Aix Marseille Université, UMR 7289 CNRS, 13005, Marseille, France

**Keywords:** Mediation analysis, local field potential, intracranial eeg

## Abstract

Neural signal shapes convey significant information about their generating processes. In this study, we introduce a data-driven methodology to identify sensory and behaviourally-relevant traces within neural responses. We present a phenomenological model that characterises temporal variations in intracranial EEG using eight interpretable parameters: peak time, peak intensity, initial and final baselines, accumulation and depletion period, and their respective concavities. This model effectively captures subtle signal variations, especially in sensory decision-making tasks. By decomposing the signals in this manner, we then conduct a comprehensive brain mediation analysis on iEEG data’s shape, pinpointing regions that mediate behavioural processes. Importantly, we can determine which signal dynamics specifically reflect underlying behavioural processes, enhancing the depth of analysis and critique of their role in behaviour. Preliminary applications on a cohort of epileptic patients reveal that our model explains over a third of the signal variance at the trial level across all brain regions. We identified four key regions—encompassing sensory, associative, frontal, and premotor areas—that mediate the impact of task difficulty on reaction time. Notably, in these regions, it was the depletion period, rather than signal amplitude, that correlated with behavioural speed. This study highlights the potential of our approach in providing detailed insights into the neural mechanisms linking stimuli to behaviour.

## Introduction

One of the key challenges in system and cognitive neuroscience is to understand how sensory information is processed by the brain and transformed to generate behaviours. Traditionally, statistical tools in neuroscience treat sensory and behavioural variables as separate. This has led to a predominant focus on correlations, often overlooking the nuanced neural processes that act as intermediaries between sensory stimuli and behavioural outcomes. Bridging this gap is essential for a more holistic understanding of how the brain encodes and generates behaviour. Recent work has proposed a new framework redefining the “neural code” as the neural features that carry sensory information used by the animal to drive appropriate behaviour (Panzeri et al., 2017). Recent metrics based on information theory have been proposed to account for the “intersection” between sensory and behavioural information (Pica et al., 2017).

Mediation analysis stands out as a simple yet structured method to probe the potential causal links between environmental stimuli, neural activity and behavioural results. It offers a way to move beyond mere observation of associations and toward a deeper understanding of the role of neural activity in cognitive processes. The limitation, however, lies in the interpretability of such results. Neural mediation tests put the nervous system as an intermediate processing step between stimulus and behaviour (Brochard & Daunizeau, 2022; Rigoli et al., 2016). Yet they can surprisingly provide little insight into neural organisations. The issue is not new and has been pointed out before, especially for neuro-imagery analysis. Many neural studies default to the analysis of overly averaged signals and artificially disconnected data points. While originally grounded on technical limitations, it risks oversimplifying neural processing, neglecting temporal dynamics and the influence of lagged responses. Complex cognitive operations, such as decision-making and choice, may require a more nuanced approach that considers, for one, temporal aspects.

In the current study, we propose a novel approach that combines mediation analysis with the variability of the neural response’s shape. Our phenomenological model allows the extraction of key features characterising neural responses, with a minimal computational burden. Mediation analysis on “shape features” allows capturing the intermediate steps that lead from stimuli to behaviour bypassing problems related to variability in sensory-motor transmission. We termed such an approach shape mediation analysis (SMA).

Previous studies have explored burst analysis and reaction times, providing valuable insights into neural and behavioural variability.These findings serve as a foundation for understanding the intricate relationship between neural responses and behaviour. SMA extends this research by offering a systematic and quantitative approach to identify intermediate factors in the neural pathway. Critically it separates behaviourally relevant features from purely physiological dynamics of the response, offering deeper insight into the underlying driver of behaviour variability. Intracranial EEG (iEEG) studies often face challenges related to statistical power due to limited data samples and noise. Data-driven approaches are essential for discovering hidden patterns, but they require robust statistical methods to avoid spurious findings. The simplicity of SMA not only addresses statistical limitations, but also enhances efficiency by reducing the number of statistical tests and enabling whole-brain exploration.

## Methods

### Neurophysiological data and experimental design

#### Stimuli and experimental design

The stimuli employed in this study were adapted from a classic visual search test originally developed by (Treisman & Gelade, 1980). Each stimulus comprised a grid of 36 letters arranged in a 6×6 square formation, with 35 “Ls” and one randomly positioned “T”. Participants were instructed to swiftly locate the “T” and press a response button upon discovery. The study featured two primary experimental conditions for comparison: an easy search condition and a challenging search condition, as illustrated in Fig. 1A. In the EASY condition, the target appeared in gray, while all distractors were in black. To distinguish between correct and incorrect responses, participants had to indicate whether the target was in the upper or lower half of the display by pressing one of two response buttons. In the DIFFICULT condition, both the distractors and the target were presented in gray. These challenging and easy condition stimuli were randomly presented for a fixed duration of 3 seconds, with a 1-second interval between each presentation. The stimuli were displayed on a 19-inch computer screen positioned 60 cm away from the participants. Each experiment was composed of 6 runs of 5-minute recording blocks. On average, participants performed 45 and 40 trials in the easy and hard conditions (Fig. 1B). They had an average of 88% and 82% of correct answers in the two conditions (Fig 1. C) with a respective response time of 1.2s ± 0.4 and 1.7s ± 0.3 (Fig. 1D).

**Fig. 1:**
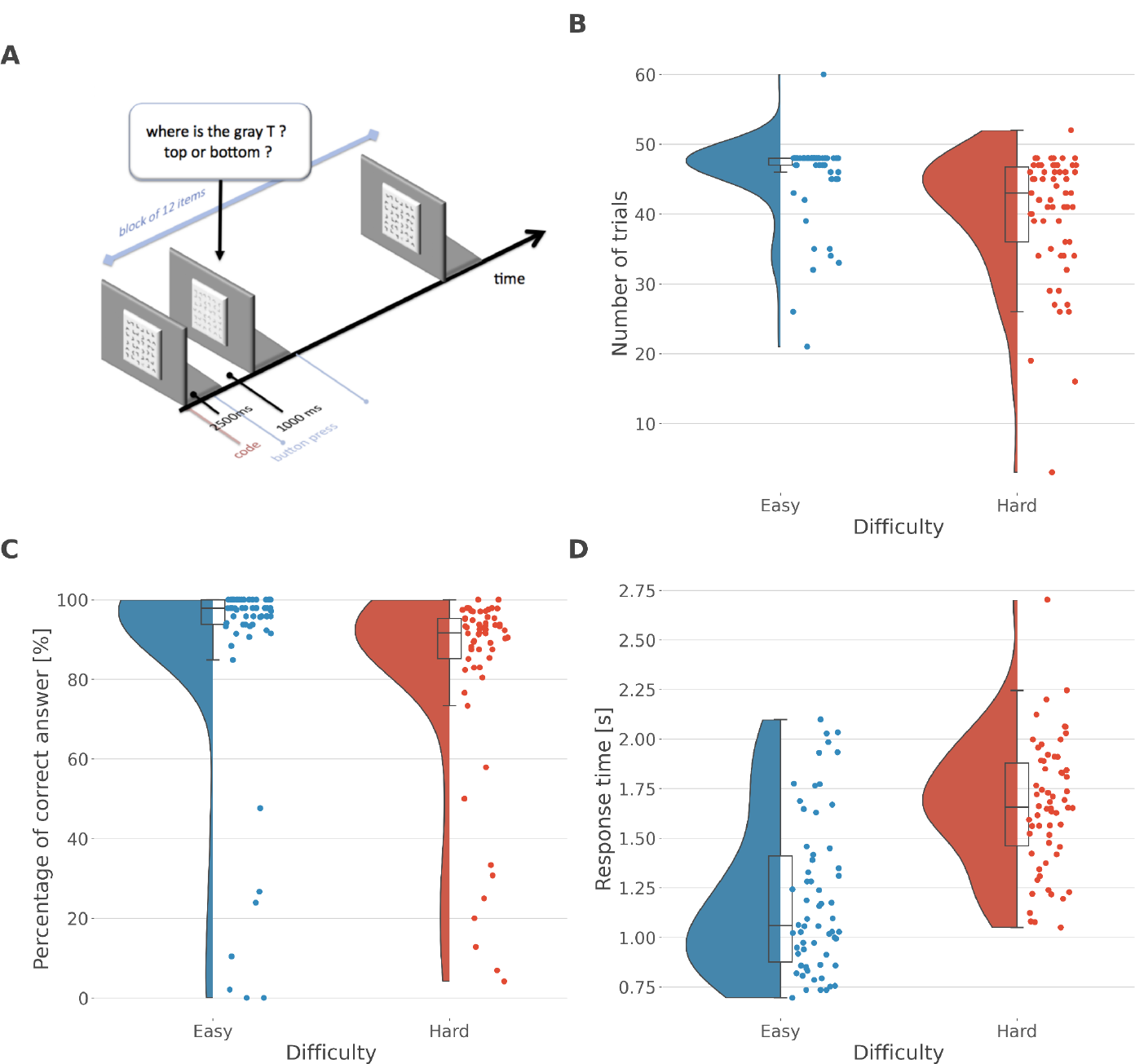
Task and behaviour. *(A)* Behavioural task; *(B)* Number of easy and hard trials across subjects, (*C*) percentage of correct answer easy and hard trials across subjects, *(D)* Response time for easy and hard trials across subjects

#### Participants

Intracranial recordings were gathered from 62 individuals undergoing neurosurgery for intractable epilepsy at the Epilepsy Department of Grenoble Neurological Hospital in Grenoble, France. All patients received stereotactic implantation of multi-lead EEG depth electrodes. Any electrode data displaying abnormal waveforms were excluded from the current study. This process involved collaboration with the medical team and entailed a visual inspection of the recordings, systematically omitting data from electrodes subsequently identified as being located within the seizure onset zone. All participants provided written informed consent, and the experimental procedures received approval from both the Institutional Review Board and the National French Science Ethical Committee.

#### Electrode implantation

Between eleven and fifteen semi-rigid multi-lead electrodes were surgically implanted in each patient using stereotactic techniques. These sEEG (stereotactic electroencephalography) electrodes had a diameter of 0.8 mm and, depending on the specific target structure, comprised 10 to 15 contact leads, each 2 mm wide and spaced 1.5 mm apart (manufactured by DIXI Medical Instruments). To ensure precise localization, all electrode contacts were first marked on the individual stereotactic implantation plan for each patient and were subsequently mapped anatomically using Talairach and Tournoux’s proportional atlas (Talairach et al., 1993). Additionally, computer-assisted alignment between a post-implantation CT scan and a pre-implantation 3-D MRI dataset (utilising VoXim R by IVS Solutions) enabled direct visualisation of the electrode contacts within the patient’s brain anatomy through ACTIVIS (developed by INSERM U1028, CERMEP, and UMR 5230). When available, a post-implantation MRI was also employed to confirm electrode positioning. The visual examination of the electrodes overlaid on each subject’s individual MRI scan was performed to ascertain whether each sEEG electrode was situated in gray or white matter. The Talairach coordinates of each electrode were finally converted into the MNI coordinate system according to standard routines (Jerbi et al., 2009, 2010; Ossandon et al., 2011; Bastin et al., 2016). Intracranial implantation and cortical repartitions of the number of recordings and subjects are displayed in Fig. 2.

**Fig. 2:**
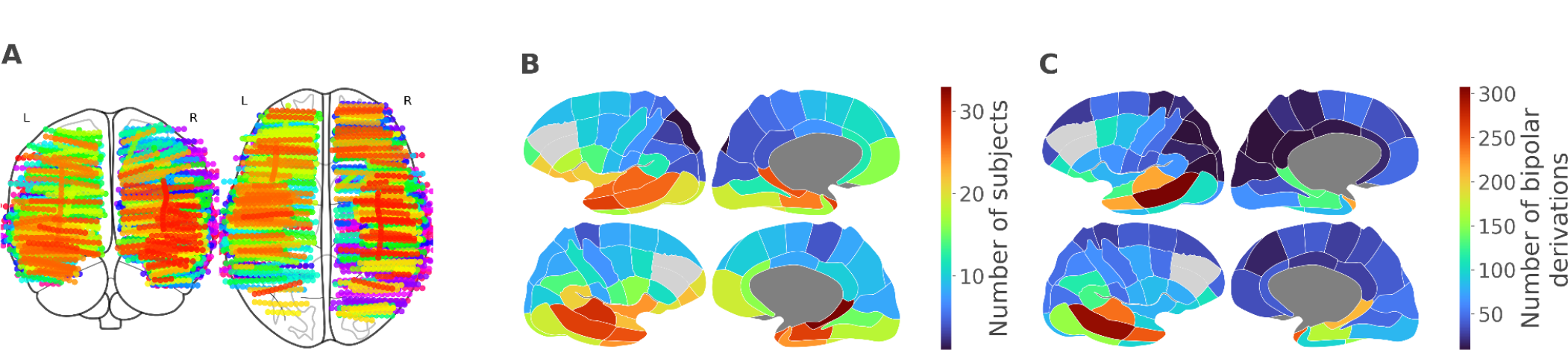
iEEG implantation and cortical repartition. *(A)* Anatomical location of intracerebral electrodes across the 62 epileptic patients. *(B)* Number of unique subjects represented per cortical parcel, *(C)* Number of of bipolar derivations per cortical parcel

#### sEEG recordings

Intracerebral recordings were conducted using a video-SEEG monitoring system (provided by Micromed), enabling simultaneous data recording from 128 depth-EEG electrode sites. The data were subjected to online bandpass filtering within the 0.1 to 200 Hz range and sampled at a rate of either 512 or 1024 Hz. During acquisition, the data were initially recorded with reference to an electrode situated in white matter. Subsequently, each electrode trace was re-referenced to its immediate neighbour, forming a bipolar derivation. This bipolar configuration offers several advantages over common referencing methods. It effectively mitigates signal artefacts commonly observed in adjacent electrode contacts (e.g., the 50 Hz mains artefact or distant physiological artefacts) and enhances local specificity by cancelling out the influences of distant sources that distribute equally to both neighbouring sites through volume conduction. The spatial resolution achieved through the bipolar SEEG configuration is approximately 3 mm, as reported in previous studies (Jerbi et al., 2009; Kahane et al., 2006; Lachaux et al., 2003). The location of all sEEG bipolar derivations was labelled according to MarsAtlas parcellation (Auzias et al., 2016).

#### Extraction of gamma envelope

To extract the gamma power envelope, we followed established procedures as outlined in prior work (Ossandón et al., 2011, 2012). In brief, we initiated the process by applying a series of continuous iEEG signal bandpass filters, spanning multiple 10-Hz-wide frequency bands. This encompassed 10 distinct bands, ranging from 50 – 60 Hz to 140 –150 Hz. Subsequently, for each signal filtered within these bands, we computed the envelope using the standard Hilbert transform. The resulting envelope possessed a time resolution of 15.625 ms. For each individual frequency band, we then normalised the envelope signal (representing time-varying amplitude) by dividing it by the mean value across the entire recording session and multiplying the result by 100. This transformation yielded instantaneous envelope values expressed as a percentage of the mean. Finally, the envelope signals computed for each consecutive frequency band (comprising 10 bands with 10 Hz intervals between 50 and 150 Hz) were amalgamated by averaging, resulting in a single time-series known as the high gamma-band envelope across the entire recording session. Notably, by design, the mean value of this time-series throughout the recording session equaled 100. Finally, the gamma envelope was down-sampled to 64 Hz. Examples of time courses of instantaneous gamma power are represented in Fig. 3.

**Fig. 3:**
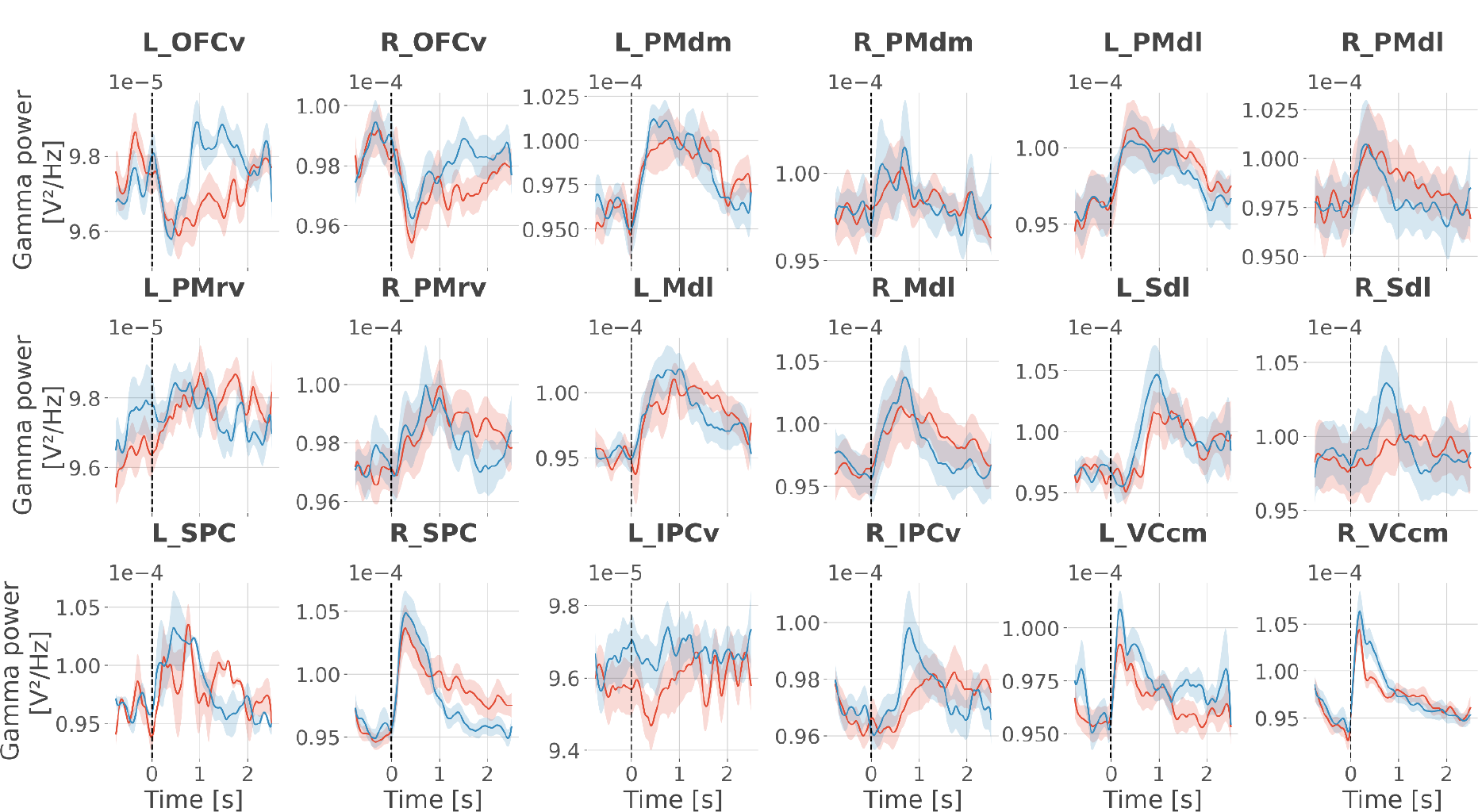
Time courses of instantaneous gamma power. The solid line and the shaded area represent the mean and SEM of the across-subjects gamma power estimated for easy trials (blue) and hard trials (red). Data are aligned to the stimulus presentation (vertical line at 0 seconds).

**Figure 4:**
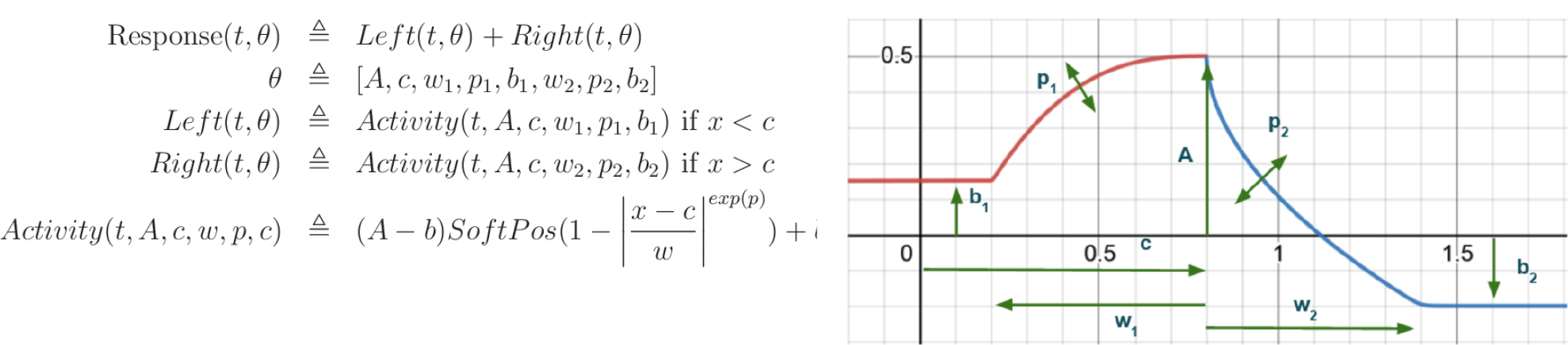
Equations governing the phenomenological model (left) and illustrations of the parameters impacts in green, on top of the red left part and the blue right part (right).

**Figure 5:**
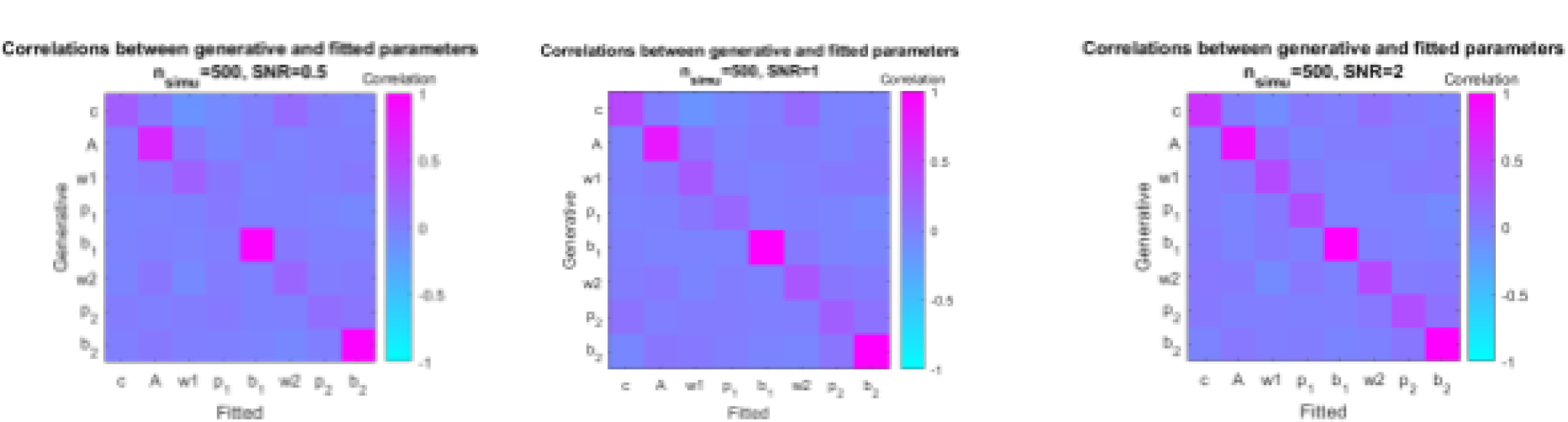
Correlation matrix of the generative and fitted parameters averaged across all trials and separated per SNR.

**Figure 6:**
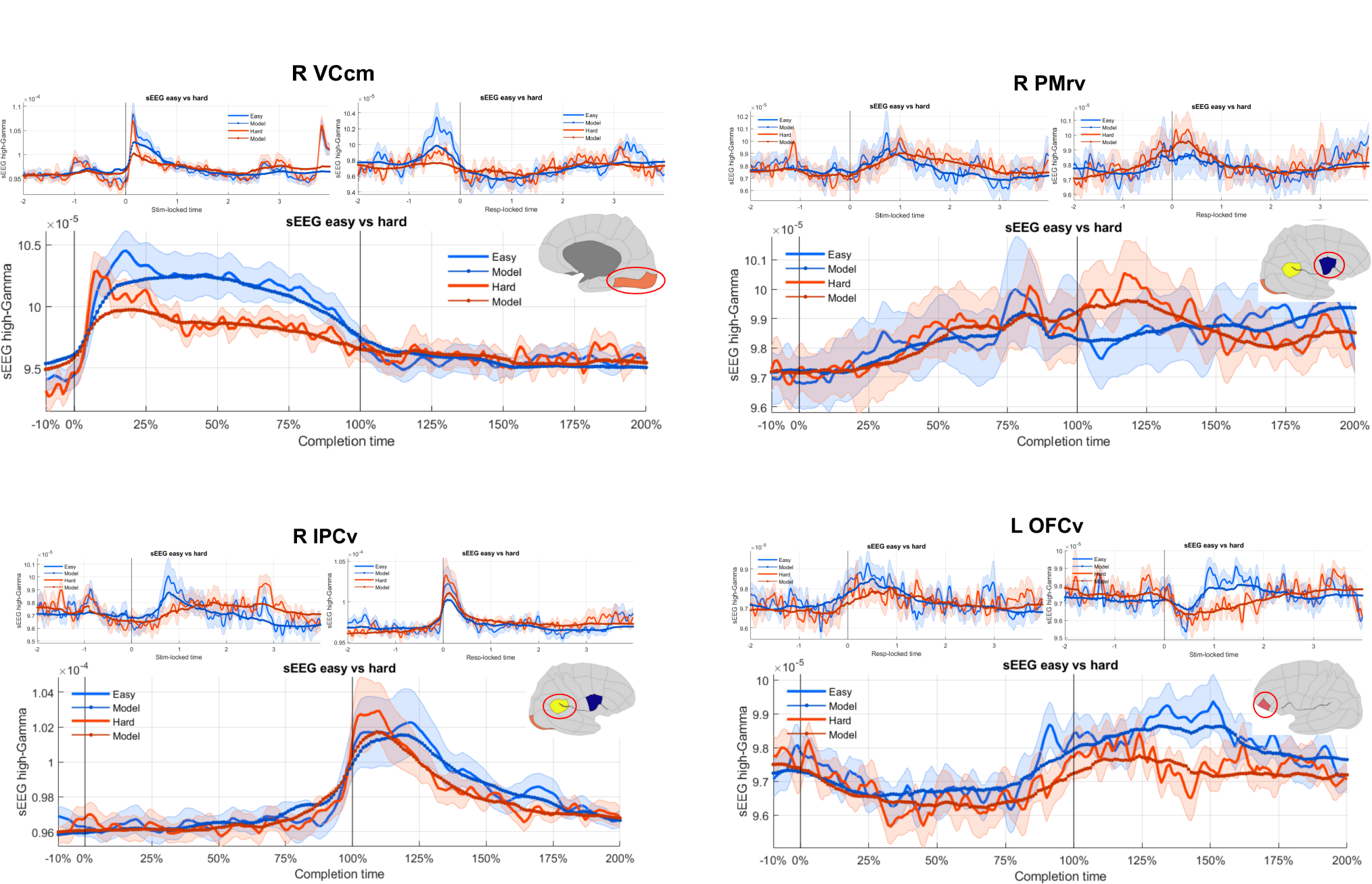
Signal variations with difficulty level (blue vs red) in the four mediating regions. All time-scale are represented for visualisation: Stimulus-locked (top left), Response-locked (top right), Completion-time (bottom). Plains curves and shaded area represent the signal’s mean and estimation error across participants, along side the mean fitted signal (dotted curves).

**Figure 7:**
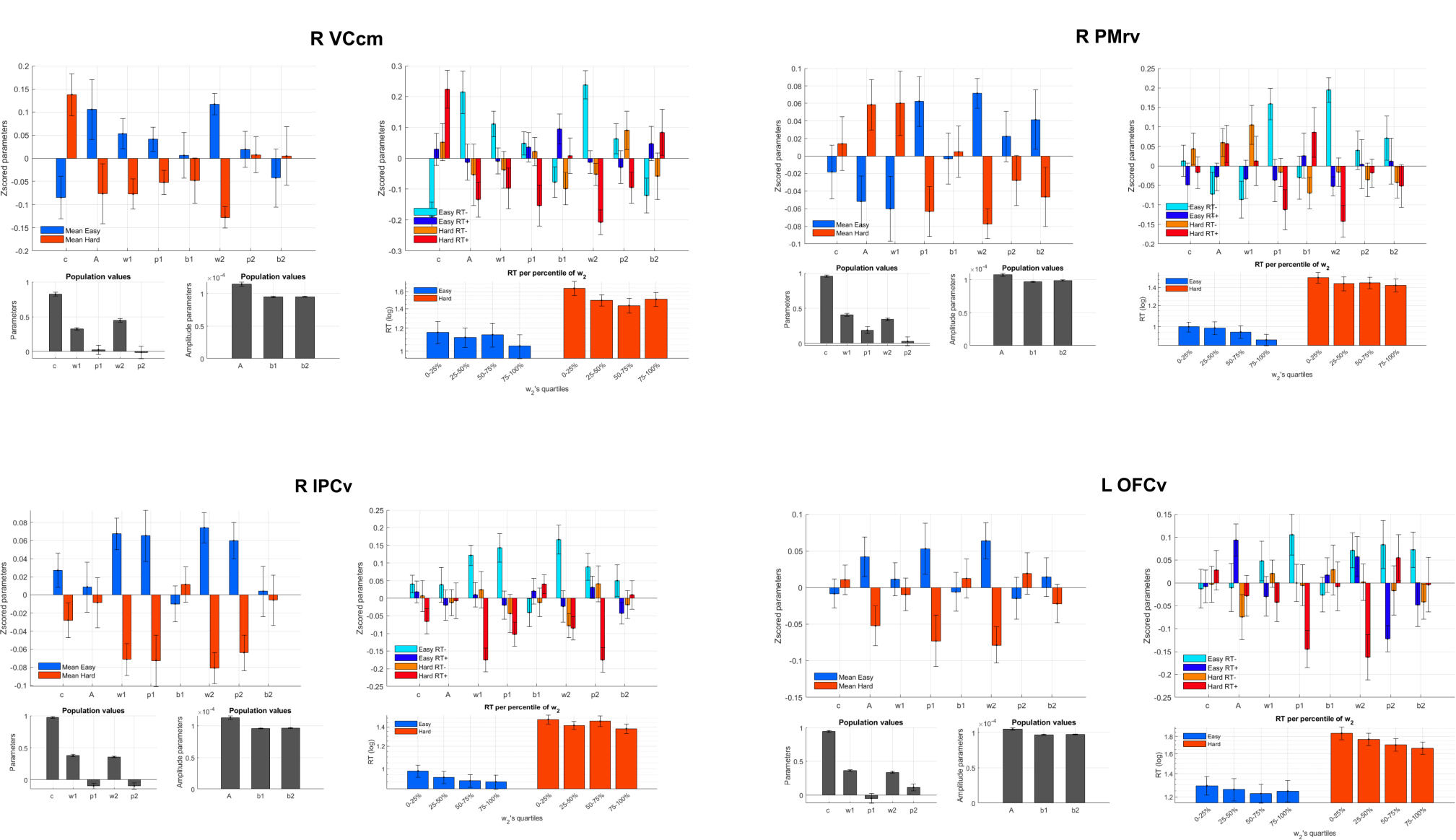
Parameters of the shape model in the four mediating regions. Upper row: separated by difficulty level (left), and response time intensity (median split per participant). Lower row: Overall value of each shape parameter (left) and relationship between participant depletion window size and reaction time (RT).

### Shape Model

In this section, we describe the construction and parameters of the shape model used to analyze neural responses.

#### Model Parameters

The central component of our analysis is a descriptive model of neural response shape described from eight simple parameters. They capture various aspects of the temporal characteristics of neural signals. Specifically:

1. **Peak Time (c):** The time point at which the neural response reaches its extrema.
2. **Amplitude (A):** Amplitude of the signal at its extrema.
3. **Initial Baseline (b1):** The baseline level preceding the response.
4. **Finishing Baseline (b2):** The baseline level following the response.
5. **Integration Time Window (w1):** The duration of the activity before its extrema.
6. **Depletion Time Window (w2):** The duration of the activity after its extrema.
7. **Concavity of Integration (p1):** The concavity of the activity before its extrema.
8. **Concavity of Depletion (p2):** The concavity of the activity after its extrema.

They are all integrated together in four simple steps to retain simple interpretability:

1. The distance to the peak is first computed and scaled: |(t-c)/w|
2. The approach dynamic of the signal is modulated by the concavity parameter |(t-c)/w|^p
3. The signal is reversed and bounded by a positivity function preventing infinite values : Pos(1-|(t-c)/w|^p)
4. Its peak and baseline are finally scaled: b + (A-b) * Pos(1-|(t-c)/w|^p)

#### Parameter Estimation

To obtain an estimate of these parameters for each electrode and each trial, we implemented a one-pass hierarchical estimation procedure. We first fitted parameters through gradient descent to the iEEG signal averaged across all levels: brain regions, participants, electrodes and experimental trials. Then we used these parameters as a starting value for the next level and rerun the fit separately for every brain region averaging the signal over the remaining levels, then again at the participants’ level, then again for each electrode and finally for each experimental trial. Ultimately, these provided a granular characterisation of the signal at multiple levels of abstraction with a relatively low computational burden.

## Statistical Analysis

In this section, we outline the statistical approaches employed to analyse the neural response data obtained through the shape model.

### Parameter recovery analysis

To validate both the shape model and our fitting procedure we randomly draw generative parameter values, generate synthetic data under various noise levels, and recover

#### parameter estimates

The correlation matrix between generative and estimated parameters captures both the recoverability of each parameter and its independence from the other.

#### Mediation Analysis

We applied mediation analysis techniques to investigate the interplay between the stimulus type, each aspect of the neural response and the subsequent behavioural outcomes. Concretely two correlation factors were evaluated across every trial, for each participant, region and electrode. The first assesses how much each parameter covaries with the difficulty level of the trials, the second how, beyond this first relation, each parameter also covaries with reaction times. A mediation test then reduces to testing the simultaneity of these relations. In this work, we focus on group-level statistics in each region. Across participants, both links need to be separately significant for a brain region to pass the test.

#### Multiple Comparison Correction

Given the large number of electrodes, brain regions and participants, we gradually reduced the number of tests to control type I errors and maintain statistical power. First, for each brain region of each subject, we average the correlation coefficients across electrodes, to retain one value per mediation path, per subject. Then each region is tested at the group level for the behavioural relationship, and the tests are corrected for all the brain regions with a FDR-correction. Finally, the selected regions are tested for the stimulus relationship and FDR corrected as well. This procedure avoids the explosion of comparison induced by the number of electrodes and avoids superfluous double tests.

## Results

### Modelling intracranial local field potential’s shape

#### Parameter recovery

To simulate synthetic signals, we used a time window of 3.2 seconds and repeatedly drew shape parameters from uniform distributions. A, b1, b2, p1 and p2, were drawn from the interval [-1,1], c from [1,2] and w1 and w2 from [0.25,1]. This ensured that all the variability of the model was represented in the synthetic data. The generated data was then corrupted with an additive noise at a signal-to-noise (SNR) ratio of 0.5, 1 or 2. For each SNR level, 500 synthetic trials were generated and separately fitted. The figure below shows the correlation between generative and fitted parameters.

The correlation matrix shows an unbiased parameters recovery, with little confusion between the parameters (all off-diagonal correlation <0.2). At high SNR, shape parameters are almost perfectly recovered. However, noise does not affect each parameter equally; the amplitude parameters A, b1 and b2 are the more robust. This is not surprising, given that each parameter has a different impact level on the signal’s variations.

#### Electrodes fit quality

To capture any traces of behavioural processing of the stimuli, we focused on the time windows covering both the stimulus presentation and the participant response. For each trial the time was then rescaled to align every stimulus presentation with time t=0 and every response with time t=1, effectively changing the time scale to a trial-completion scale: *t*_*completion*_ < −(*t* − *t*_*stimulus*_)/*RT* . This simple procedure allowed us to examine both types of sensitivity without artificially biasing the models with variable time windows. In this setting, our model explained on average 33% (std 5.25%) of signal variances across all ROIs. Respectively 37% (std 5.5%) and 30% (std 5%) for easy and hard trials. This quality of fit decreased to 22% (std 4.3%) and 22% (std 4.2%) when aligning the signals to either the stimulus or the response.

**Table 1:**
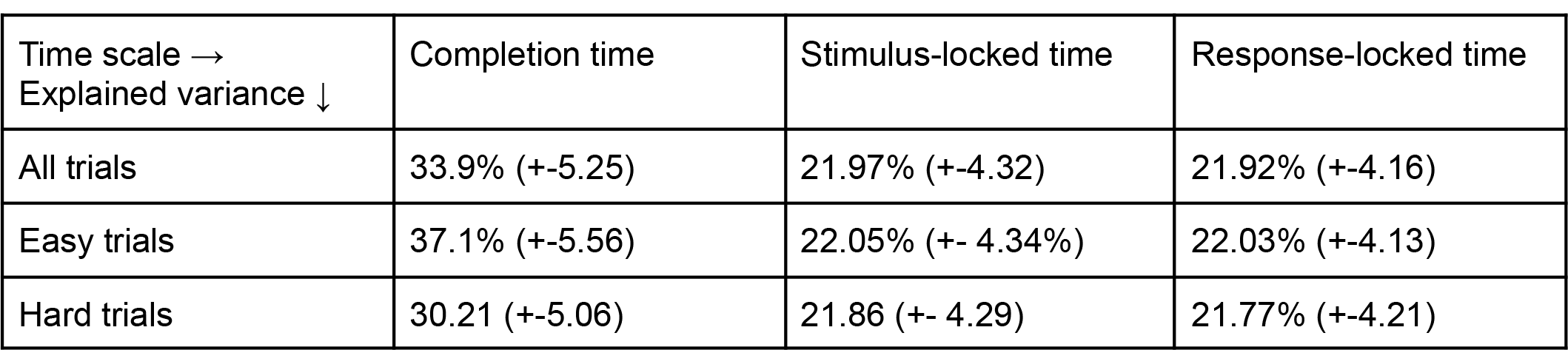
Fit quality across the Completion-time scale, the Stimulus-locked time scale and the Response-locked time scale.

### Mediation analysis on neural shape parameters

#### Mediation analysis

After FDR correction across the 82 regions, four regions yield significant mediation results: the right caudal medial Visual Cortex (R VCcm), rostral ventral Premotor Cortex (R PMrv), ventral Inferior Parietal Cortex (R IPCv) and the left ventral Orbito Frontal Cortex (L OFCv). Overall, the model explained 38% of their electrodes’ variance. All four regions mediated the effect of the task difficulty on the reaction time through the length of the depletion time window (w2). Switching from easy to hard trials increased each region’s depletion window, relative to the reaction time. In turn, increases in w2 were associated with a shortening of the reaction time, above and beyond the changes induced by the difficulty level.

**Table 2:**
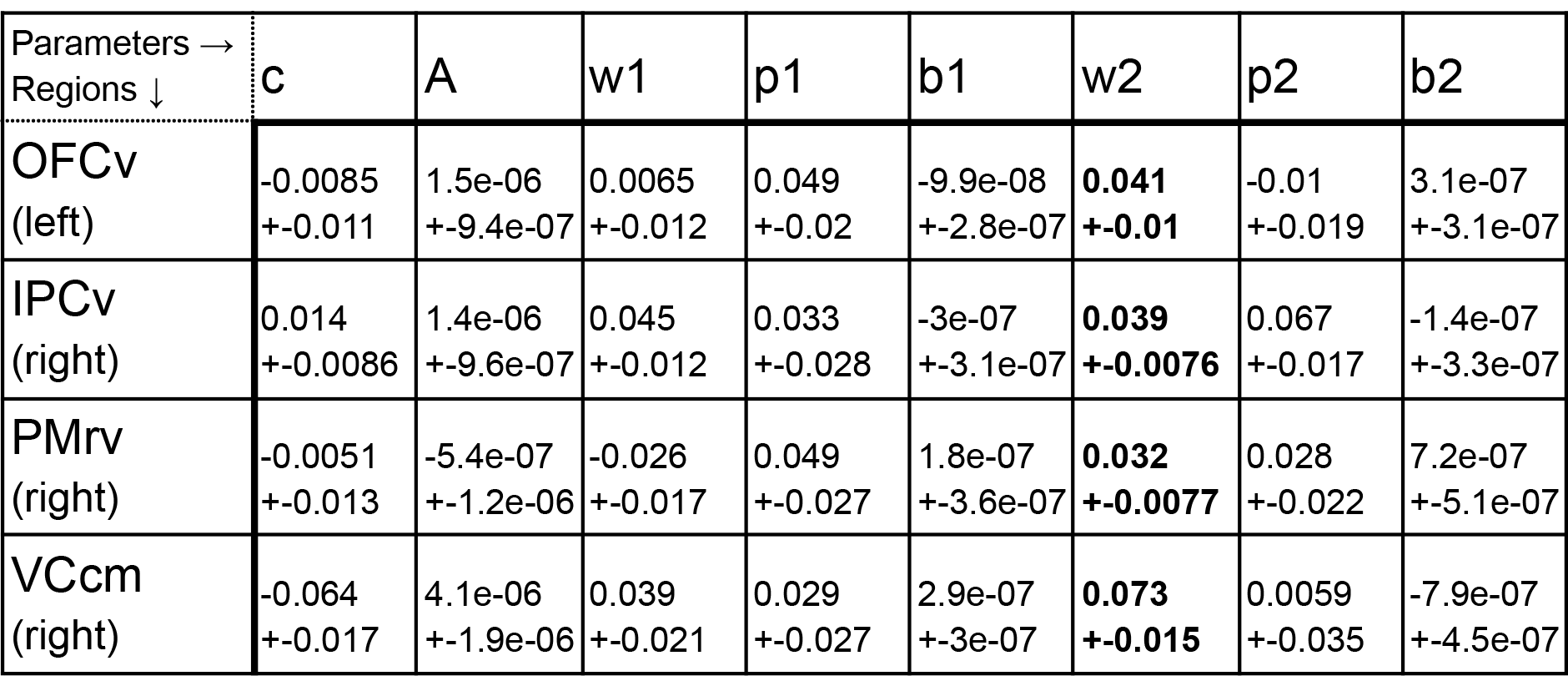
Strength of the mediation’s path between the task difficulty and each shape parameter, in each region alongside their estimation’s error. Significant elements (FDR-corrected) are bolded.

**Table 3:**
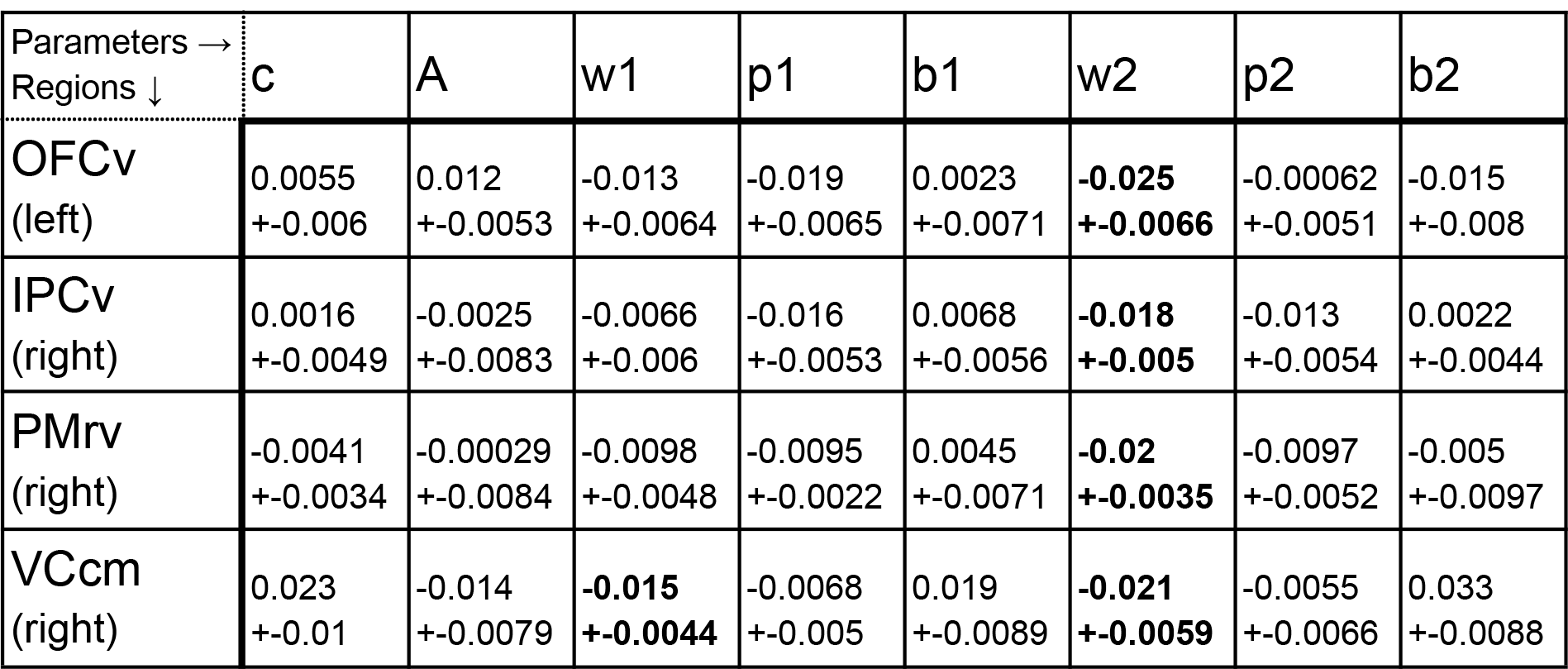
Strength of the mediation’s path between each shape parameter and the subject’s reaction time, in each region alongside their estimation’s error. Significant elements (FDR-corrected) are bolded.

## Discussion

In our study, we aimed to identify neural processes related to the processing of sensory information that leads to behavioural control. To achieve this, we employed a novel technique, the Shape Mediation Analysis (SMA), to gain deeper insights into the temporal patterns present in iEEG signals. Our approach decomposes these signals into eight distinct features, allowing us to observe their variations across different trials. This detailed analysis enabled us to pinpoint features that simultaneously reflect sensory sensitivity and describe behaviour.

SMA’s application to iEEG effectively bridges the detailed physiological recording of iEEG toward the broad discovery potential seen in fMRI studies. However, instead of relying solely on extensive activation maps, SMA emphasizes a select set of features, making it conducive to modelling neural populations and their interconnections. Our findings underscore that focusing on neural activation limits results by weakening statistical sensitivity. By examining other signal features, we can obtain a richer understanding of how different neural populations interact across brain regions.

Our results not only corroborate but also refine disparate findings from the neuro-imaging literature on decision-making. For instance, the involvement of the VCcm in attentional and risk evaluation processes is supported by works like (Giesbrecht et al., 2006; Krain et al., 2006) . The IPCv’s association with motor selection, action evaluation, evidence accumulation, and exploration in studies by (Daw et al., 2006; Gluth et al., 2012; Krueger et al., 2017; Madlon-Kay et al., 2013; Paulus et al., 2005). The PMrv, highlighted in many of these studies, is not only linked to information gathering and uncertainty evaluation (as seen in (Krug et al., 2014)) but also showcases a functional organization spanning accumulation, selection, and execution(Rebola et al., 2012). This suggests a need for a more detailed segmentation than what we used with marsAtlas. Lastly, the OFCv’s role in valuing decisions under uncertainty (Krug et al., 2014) and risky behaviour (Fishbein et al., 2005) further complement our findings. Our approach binds all these results within one interpretable data-driven framework.

## References

Auzias, G., Coulon, O., & Brovelli, A. (2016). MarsAtlas : A cortical parcellation atlas for functional mapping. Human Brain Mapping, 37(4), 1573–1592. 10.1002/hbm.23121

Brochard, J., & Daunizeau, J. (2022). Synaptic plasticity in the orbitofrontal cortex explains how risk attitude adapts to the range of risk prospects (p. 2020.09.08.287714). bioRxiv. 10.1101/2020.09.08.287714

Daw, N. D., O’Doherty, J. P., Dayan, P., Seymour, B., & Dolan, R. J. (2006). Cortical substrates for exploratory decisions in humans. Nature, 441(7095), Article 7095. 10.1038/nature04766

Fishbein, D. H., Eldreth, D. L., Hyde, C., Matochik, J. A., London, E. D., Contoreggi, C., Kurian, V., Kimes, A. S., Breeden, A., & Grant, S. (2005). Risky decision making and the anterior cingulate cortex in abstinent drug abusers and nonusers. Brain Research. Cognitive Brain Research, 23(1), 119–136. 10.1016/j.cogbrainres.2004.12.010

Giesbrecht, B., Weissman, D. H., Woldorff, M. G., & Mangun, G. R. (2006). Pre-target activity in visual cortex predicts behavioral performance on spatial and feature attention tasks. Brain Research, 1080(1), 63–72. 10.1016/j.brainres.2005.09.068

Gluth, S., Rieskamp, J., & Büchel, C. (2012). Deciding When to Decide : Time-Variant Sequential Sampling Models Explain the Emergence of Value-Based Decisions in the Human Brain. Journal of Neuroscience, 32(31), 10686–10698. 10.1523/JNEUROSCI.0727-12.2012

Jerbi, K., Ossandón, T., Hamamé, C. M., Senova, S., Dalal, S. S., Jung, J., Minotti, L., Bertrand, O.,Berthoz, A., & Kahane, P. (2009). Task-related gamma-band dynamics from an intracerebral perspective : Review and implications for surface EEG and MEG. Human brain mapping, 30(6), 1758–1771.

Kahane, P., Landré, E., Minotti, L., Francione, S., & Ryvlin, P. (2006). The Bancaud and Talairach view on the epileptogenic zone : A working hypothesis. Epileptic Disorders: International Epilepsy Journal with Videotape, 8 Suppl 2, S16–26.

Krain, A. L., Wilson, A. M., Arbuckle, R., Castellanos, F. X., & Milham, M. P. (2006). Distinct neural mechanisms of risk and ambiguity : A meta-analysis of decision-making. NeuroImage, 32(1), 477–484. 10.1016/j.neuroimage.2006.02.047

Krueger, P. M., van Vugt, M. K., Simen, P., Nystrom, L., Holmes, P., & Cohen, J. D. (2017). Evidence accumulation detected in BOLD signal using slow perceptual decision making. Journal of Neuroscience Methods, 281, 21–32. 10.1016/j.jneumeth.2017.01.012

Krug, A., Cabanis, M., Pyka, M., Pauly, K., Walter, H., Landsberg, M., Shah, N. J., Winterer, G., Wölwer, W., Musso, F., Müller, B. W., Wiedemann, G., Herrlich, J., Schnell, K., Vogeley, K., Schilbach, L., Langohr, K., Rapp, A., Klingberg, S., & Kircher, T. (2014). Investigation of decision-making under uncertainty in healthy subjects : A multi-centric fMRI study. Behavioural Brain Research, 261, 89–96. 10.1016/j.bbr.2013.12.013

Lachaux, J. P., Rudrauf, D., & Kahane, P. (2003). Intracranial EEG and human brain mapping. Journal of Physiology, Paris, 97(4-6), 613–628. 10.1016/j.jphysparis.2004.01.018

Madlon-Kay, S., Pesaran, B., & Daw, N. D. (2013). Action selection in multi-effector decision making. NeuroImage, 70, 66–79. 10.1016/j.neuroimage.2012.12.001

Ossandón, T., Jerbi, K., Vidal, J. R., Bayle, D. J., Henaff, M.-A., Jung, J., Minotti, L., Bertrand, O., Kahane, P., & Lachaux, J.-P. (2011). Transient Suppression of Broadband Gamma Power in the Default-Mode Network Is Correlated with Task Complexity and Subject Performance. Journal of Neuroscience, 31(41), 14521–14530. 10.1523/JNEUROSCI.2483-11.2011

Ossandón, T., Vidal, J. R., Ciumas, C., Jerbi, K., Hamamé, C. M., Dalal, S. S., Bertrand, O., Minotti, L., Kahane, P., & Lachaux, J.-P. (2012). Efficient “Pop-Out” Visual Search Elicits Sustained Broadband Gamma Activity in the Dorsal Attention Network. Journal of Neuroscience, 32(10), 3414–3421. 10.1523/JNEUROSCI.6048-11.2012

Panzeri, S., Harvey, C. D., Piasini, E., Latham, P. E., & Fellin, T. (2017). Cracking the Neural Code for Sensory Perception by Combining Statistics, Intervention, and Behavior. Neuron, 93(3), 491–507. 10.1016/j.neuron.2016.12.036

Paulus, M. P., Feinstein, J. S., Leland, D., & Simmons, A. N. (2005). Superior temporal gyrus and insula provide response and outcome-dependent information during assessment and action selection in a decision-making situation. NeuroImage, 25(2), 607–615. 10.1016/j.neuroimage.2004.12.055

Pica, G., Piasini, E., Safaai, H., Runyan, C., Harvey, C., Diamond, M., Kayser, C., Fellin, T., & Panzeri, S. (2017). Quantifying how much sensory information in a neural code is relevant for behavior. Advances in Neural Information Processing Systems, 30. https://proceedings.neurips.cc/paper_files/paper/2017/hash/a9813e9550fee3110373c21fa012eee7-Abstract.html

Rebola, J., Castelhano, J., Ferreira, C., & Castelo-Branco, M. (2012). Functional parcellation of the operculo-insular cortex in perceptual decision making : An fMRI study. Neuropsychologia, 50(14), 3693–3701. 10.1016/j.neuropsychologia.2012.06.020

Rigoli, F., Friston, K. J., & Dolan, R. J. (2016). Neural processes mediating contextual influences on human choice behaviour. Nature Communications, 7(1), Article 1. 10.1038/ncomms12416

Talairach, J., Tournoux, P., & Missir, O. (1993). Referentially oriented cerebral MRI anatomy : An atlas of stereotaxic anatomical correlations for gray and white matter. https://www.semanticscholar.org/paper/Referentially-oriented-cerebral-MRI-anatomy-%3A-an-of-Talairach-Tournoux/1ae2e3fcc68c38fc01486999514ed4847737f8f9

Treisman, A. M., & Gelade, G. (1980). A feature-integration theory of attention. Cognitive Psychology, 12(1), 97–136. 10.1016/0010-0285(80)90005-5

